# Meta-analysis of the variance ratio

**DOI:** 10.1101/104695

**Authors:** Nicolas Traut

**Affiliations:** Unité Génétique Humaine et Fonctions Cognitives, Département de Neuroscience, Institut Pasteur, Paris, France; Sorbonne Universités, UPMC, CNRS, Inserm, IBPS, Neuroscience Paris Seine, France

## Abstract

The most commonly used effect size when using meta-analysis to compare a measurement of interest in two different populations is the standardised mean difference. This is the mean difference of the measurement divided by the pooled standard deviation in the two groups. The standard deviation is usually supposed to be the same for both groups, although this assumption is often made without any particular evidence. It is possible, however, that the difference of the measurement in the two populations resides precisely in their standard deviations. This could be the case, for example, if a population of patients exhibited more “abnormal” values than a control population – both large and small – even if the mean values were the same. Fisher’s test of equality of variance is designed to compare standard deviations. A variance ratio is a Fisher’s ratio and Fisher distribution can be used to give confidence intervals to the estimate for one study. However, confidence interval for one study can be very wide if the study does not contain enough subjects. Here we present an approach to combine variance ratios of different studies in a meta-analytic way which produces more robust estimates under these circumstances.

## Method

The problem of combining directly variance ratio effect sizes is that the the Fisher distribution is far from the normal distribution. Meta-analyses using fixed effect models or random effects models are designed to combine effect sizes obtained from samples following a normal distribution. Therefore, we prefer to consider the log-variance ratio as effect size 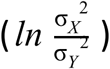 because the distribution of its estimator is approximately normal. In what follows we will show that an unbiased estimator for the log-variance ratio is

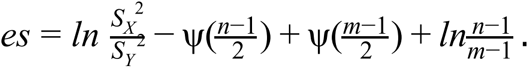

We begin by looking at the natural estimator of 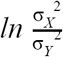:

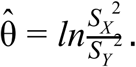

The distribution of this estimator can be evaluated using the following development, where n is the size of sample X and m the size of sample Y:

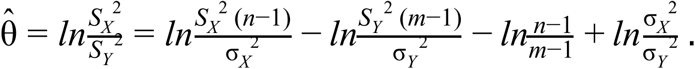

The application of Cochran's theorem tells us that if X and Y are distributed according to a normal distribution, variables 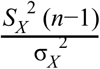 and 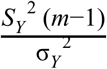 are each distributed according to a *χ*^2^ distribution of n − 1 et m − 1 degrees of freedom respectively. The estimator of the effect is therefore distributed as a linear combination of variables following the logarithm of a *χ*^2^ distribution. It has been shown that the *χ*^2^ distribution is well approximated by a lognormal distribution (Jouini et al. 2011), therefore the distribution of the estimator can be effectively approximated as being normal.

To use the estimator in the meta-analysis, we need to know its expectation and its variance. One can show for *Q* ~ *χ*^2^(*ν*) that:

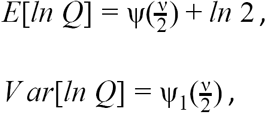

where ψ(*x*) is the digamma function, and ψ_1_ (*x*) is the trigamma function.

We deduce that

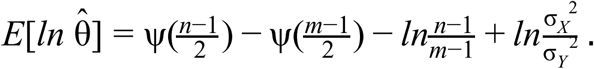

The estimator is consequently biased, so we replace it by the following unbiased estimator:

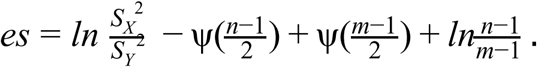

The expectation of the new estimator is 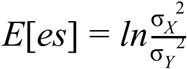 and its variance is 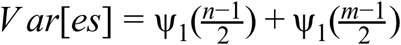.

## Simulations

We simulated an example of meta-analysis of 10 studies where the distributions of the value of interest for cases and controls were the same but the standard deviation for cases was 1.2 times larger than the standard deviation for controls. For the illustration of the impact of bias removing in the estimation of the log-variance ratio, we made different sample sizes for cases and controls: sample sizes were derived from an uniform random distribution ranging from 20 to 40 for cases and from an uniform random distribution ranging from 5 to 30 for controls. We assessed the validity and the precision of the estimations by doing a Monte Carlo simulation with 10,000 independent meta-analyses. We evaluated the existence of a bias in the meta-analysis by comparing the set of estimated log-variance ratios to the real value with a z-test. The R code for doing the simulations was written in a Jupyter notebook and could be accessed here: https://github.com/ntraut/My-notebooks/blob/master/lvr-meta.ipynb.

## Results

As specified in the simulation parameters, the mean total number of cases was larger than the mean total number of controls (300 vs 175). As a result, the log-variance ratio was slightly overestimated without bias correction (p<10^−100^): with our parameters, the mean of estimations was 0.394 when the real value was 0.365. At the opposite, the correction for the bias seemed to work (in our run, p=0.107 for the fixed effect model and p=0.408 for the random effects model). The statistical power for the meta-analysis was 69.4% for the fixed effect model, with an error rate of 5.07%, which is comparable to the chosen alpha level of 5%. In our case, the random effects model was overly conservative with a error rate of 3.84%. This can be understood by the fact that in our case, studies were homogeneous by design, so the fixed effect model would have been the appropriate model. The error rate was not considerably increased when not correcting for bias in the estimation of log-variance ratio: it was 5.36% in fixed effect model and 4.06% in random effects model. The results of the Monte Carlo simulation are also described in **Table 1**.

**Table 1.** Monte Carlo simulation results SD: standard deviation; SEM: standard error of the mean; p-value: probability for obtaining a mean of estimations at least as far from the real value as the observed difference, used to assess for the existence of a bias in the estimations; statistical power: estimated probability for obtaining a combined log-variance ratio significantly different from 0; frequency incorrect: rate of times where estimated 95% intervals did not overlap the real value.

|  | Model | Distribution of estimations (real value: 0.3646) |  |  |  | Statistical power | Frequency incorrect |
| --- | --- | --- | --- | --- | --- | --- | --- |
|  |  | Mean | SD | SEM | p-value |  |  |
| With bias correction | Fixed effect | 0.3623 | 0.1470 | 0.0015 | 0.107 | 69.4% | 5.07% |
|  | Random effects | 0.3634 | 0.1481 | 0.0015 | 0.407 | 62.9% | 3.84% |
| Without bias correction | Fixed effect | 0.3939 | 0.1474 | 0.0015 | 0 | 76.2% | 5.36% |
|  | Random effects | 0.3974 | 0.1490 | 0.0015 | 0 | 70.8% | 4.06% |

## Discussion

We presented a method to compare standard deviations of two populations in a meta-analysis. The Monte Carlo simulation showed that this method allowed us to obtain a reasonable statistical power to detect relatively small difference of standard deviations: in our case, the statistical power was 69% to detect a ratio of standard deviations of 1.2 with 300 cases and 175 cased split in 10 studies. This statistical power was slightly below than the statistical power that we would have obtained if we had made a single F-test on the merged samples (76%), but the meta-analytical approach allows to take account of a possible study effect, for example when all the studies do not use exactly the same scale or are heterogeneous. The Monte Carlo simulation did not show evidence of a bias when we used the corrected log-variance ratio, but the uncorrected log-variance ratio was clearly biased, although the differences of estimation were relatively small.

This type of meta-analysis needs to be tested in other meta-analyses in order to evaluate its capacity to find statistically significant results when possibly the meta-analysis on mean differences does not find any. We ourselves used this method in a meta-analysis on cerebellar volume alterations in autism, which should shortly be published.

